# Predicting Antifouling Paint Particle Contamination based on 16S rRNA Gene Sequencing Data using Random Forest-Based Machine Learning

**DOI:** 10.1101/2025.09.03.673959

**Authors:** T. Sperlea, M Labrenz, B. Kreikemeyer, A. Schenk, A. S. Tagg

**Affiliations:** Leibniz Institute for Baltic Sea Research Warnemünde, Rostock 18119, Germany; Institute of Medical Microbiology, Virology and Hygiene, University of Rostock, 18057 Rostock, Germany

## Abstract

Antifouling paints often contain biocides designed to inhibit biological growth, and antifouling paint particles (APPs) have been previously shown to affect microbial communities in sediment. Given typical methods for monitoring for APP presence can be specialised and challenging, alternative methods using simple, standardised and universal approaches, such as 16S rRNA amplicon sequencing, would be highly valuable. This study uses a field-based mesocosm approach to train a random forest-based (supervised) machine learning model to predict APPs presence and concentration in sediment based on 16S microbial community data. The model correctly predicted 100% APP-presence samples and 83.3% APP-absence samples in the incubation testset, although the model could not correctly predict APP concentration with sufficient accuracy. To determine real-world applicability of the model, samples from 14 different sites along the Baltic Sea coastline and Warnow estuary in NE Germany were collected and APP-presence was pre-determined using SEM-EDX spectroscopy. The model correctly assigned APP-absence status to all APP-absent sites, and correctly assigned 3 of 5 APP-contaminated sites as having APP presence. As such, these results confirm it is possible to predict APPs in sediment based on the microbial community, and serves as a proof-of-concept for the further development of machine learning-based predictive tools for environmental monitoring.

## Introduction

Paint particles represent an important but often overlooked class of aquatic pollutants that make up an appreciable proportion of microplastics, especially in estuarine and coastal sediments.^1,2^ What distinguishes paint particles from other “typical” microplastics which are comprised almost wholly of chains of synthetic polymers such as polyethylene (PE), polypropylene (PP), polyamides (PA, including nylons) and polyethylene terephthalate (PET), is their much larger proportion of non-plastic components such as solvents, pigments and other additives such as flame retardants, binders and heavy metals.^1,3^ Antifouling paints are a subgroup of paints distinguished by the engineered ability to inhibit biological growth and biofilm formation on surfaces. Broadly, there are two subcategories of antifouling paints: hard antifouling and soft or ablative antifouling paints.^4^ Both types function through controlled release of biocidal agents from the painted surface but differ in the delivery method due to different use cases of the painted structure. Hard antifouling paints are used for stationary structures or for boats which spend a considerable time sitting stationary in water, especially freshwater. In contrast, soft antifouling paints are used for commercial boats which are frequently in motion. As the boat moves through water, the soft layers slowly and deliberately erode, exposing fresh layers of biocide over time. While both antifouling paint classes require reapplication, hard antifouling paints may present a larger concern regarding the release of antifouling paint particles (APPs) into sediment because, before they can be applied, they require a greater degree of surface preparation to remove previous material build-up. Such hull-blasting and mechanical abrasion processes are likely to contribute considerably to the load of microplastic-sized APPs in boatyards, marinas, and maintenance facilities located near estuarine and coastal environments.^5,6^ Given such loads of APPs reaching environmental sediments, it is reasonable to expect particulates with slow-release biocides to have an environmental impact of sediment biology. Recent research by the authors demonstrated in a microcosm experiment that APPs, regardless of their chemical composition, induce a clear and consistent shift in the microbial community composition of coastal marine sediment, while non-antifouling paint particles caused no observable change compared to controls or the starting community.^7^ These findings highlight the need to distinguish APPs from non-antifouling paint particles in environmental monitoring efforts. However, while the environmental monitoring of antifouling paint particles is theoretically covered under general microplastic monitoring, this is often not the case due to the complexities in identifying antifouling paints compared to typical microplastics.

Identifying microplastics in environmental samples, especially at sub-500 µm sizes is already extremely labour intensive. Typically rounds of density separation and organic digestion are needed to process samples to the point where microplastics are retained for analysis while sediment and organic contaminants (i.e. biological material) have been removed.^11,12^ Only then can analysis begin where, in best-of-practise scenarios, transmission Fourier-transform infrared spectroscopy (FTIR) and Raman spectroscopy are used in combination to identify and categorise microplastics from other particulates (i.e. black carbon) which can still be retained in samples despite the extensive processing steps. This dual spectroscopic approach is particularly necessary to identify paints from typical microplastics, since although typical plastics often also contain pigments and additives, these are usually at trace levels. In contrast, non-bulk polymer components can make up half or more of the total mass of paints.^1,3^ However, from an environmental perspective, perhaps the most important identification distinction in microplastics is determining whether detected paint particles have antifouling components; however, even well-developed microplastic identification analyses pipelines using FTIR and Raman spectroscopy are typically unable to identify antifouling distinctions.^8^ Instead, a third round of elemental-focused spectroscopy, such as x-ray photoelectron spectroscopy, is typically required to detect the presence of heavy metals which are particularly important for a toxicological assessment. However, adding the requirement for an additional round of complex spectroscopy (which also requires coating samples in conductive material, such as gold or carbon, which would disrupt the ability for downstream FTIR/Raman) is likely a step too far for most microplastic researchers. Indeed, despite the rapid expansion of global microplastic research over the past two decades, most surveys have been unable to determine if antifouling paint particles have been detected. This represents a glaring gap in the current understanding of microplastic pollution as APPs — unlike most inert microplastic particles — can exert a marked effect on biological processes and microbial community dynamics.^7,9,10^ Thus, the need for a better way to determine APP presence is greatly needed, in addition to the need to also further examine to what extent APPs are actually causing an effect on sediment microbiology in the environment.

The analysis of microbial communities may be able to achieve both simultaneously. The use of indicator taxa – a taxon or group of taxa which can, by their presence or activity, represent specific environmental conditions – is well-established in ecological research.^13^ Prominent examples include the marsh periwinkle (*Littoraria irrorata*), which was an important indicator for the wide-reaching effects of the Deepwater Horizon oil spill, as changes in its growth, population and productivity provided critical insights into the spill’s ecological consequences and geographical area of effect.^14^ Microbial communities are highly sensitive to environmental perturbations and can, therefore, serve as powerful bioindicators.^15,16^ Advances in high-throughput sequencing combined with decreasing cost of 16S ribosomal RNA sequencing have facilitated the routine characterization of microbial communities. Sediment environments in particular harbour huge microbial diversity compromising many thousands of genetically distinct bacterial taxa.^17^ As a result, even small environmental changes can have wide reaching effects to the microbial community composition. More recently, machine learning methods have emerged that promise to resolve subtle yet consistent community-level shifts attributable to environmental signals. These are characterized by the ability to capture nonlinear relationships in high-dimensional datasets.^18,19^ Broadly speaking, in a training phase, supervised machine learning models are exposed to microbial community composition data and the corresponding values of the variable the model should learn to approximate. The quality of the model can then be evaluated in a testing phase where the model is faced with data withheld from training in terms of the deviation between predicted and true values. Indeed, machine learning methods like Random Forests have been used to identify microbial correlates of anthropogenic interaction with the environment, such as contamination with glyphosate,^20,21^ the munition compound 2,4,6-trinitrotoluene^22^ and land use.^16^

This study has adopted a two-stage approach to assess whether the sediment microbial community composition can serve as a bioindicator for APP contamination. The first stage utilised an experimental mesocosm set-up using sediment chambers with a range of known APP contamination levels placed in Baltic Sea coastal sediments for 60 days. Following exposure, the microbial communities within these sediments were characterised using high-throughput 16S rRNA gene amplicon sequencing. This data was then used to train a Random Forest model to determine whether the level of APP contamination can be approximated based on shifts in community composition. In the second stage, the trained model was applied to a field-based survey of sediments collected from multiple locations along the German Baltic Sea coastline, including the Warnow estuary; areas within which can be characterised by intense anthropogenic maritime activity. To corroborate the model predictions, sediment samples underwent parallel EDX analyses to directly identify APPs and quantify contamination to ultimately produce a validated predictive model for future APP monitoring. If effective, the novel use of a small but directed mesocosm setup, as opposed to large-scale environmental surveying, to train a machine learning model also lays a blueprint for a new, straight-forward and inexpensive way to construct a biomonitoring predictive model for a wide range of other anthropogenic stressors.

## Results

### Mesocosm experiments suggest difference between APP-contaminated and APP-uncontaminated sediment samples

In order to capture the effect of the presence of APPs on the microbial community composition, we performed a mesocosm experiment exposing sediment samples to different levels of APPs. A Principal Coordinate Analysis-based ordination of the samples taken after 60 days of exposure in Baltic Sea coastal sediment (see Methods) demonstrates that control chamber samples (without APP addition) form a closely ordinated cluster along PCoA 1. In contrast, APP-contaminated samples exhibited substantially greater variability in their community structure. Notably, APP concentration does not appear to have a consistent, dose-dependent effect on the microbial community composition, as samples exposed to low, medium and high APP concentrations demonstrate similar variability in microbial community (see Figure 2A). Thus, the results can broadly be considered as showing only two distinctive groups: APP-contaminated (low, medium and high) and APP-uncontaminated (control) samples.

**Fig. 1.**
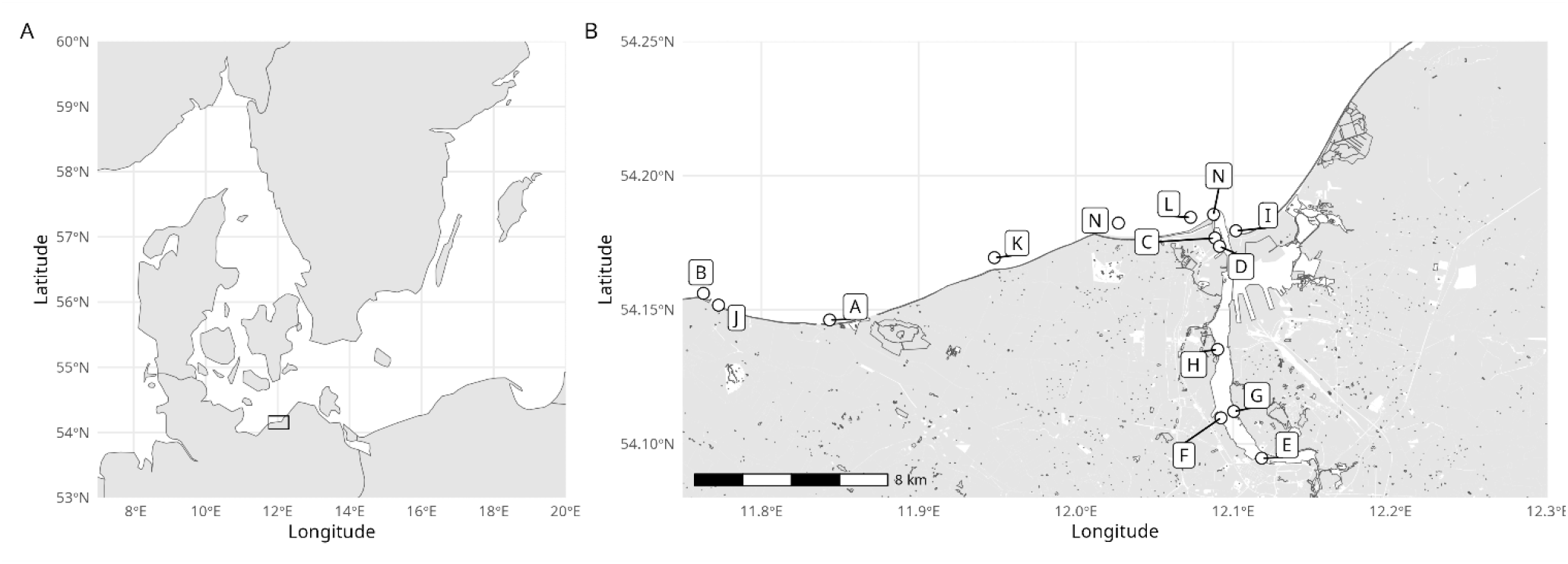
A: Image of the southern Baltic Sea. Black box highlighted the area detailed in B: Sites A - O denoting each of the individual sites where sediment was sampled both for DNA extraction and APPs for model validation. Site A also represents the location where the mesocosm experiment was conducted.

**Fig. 2.**
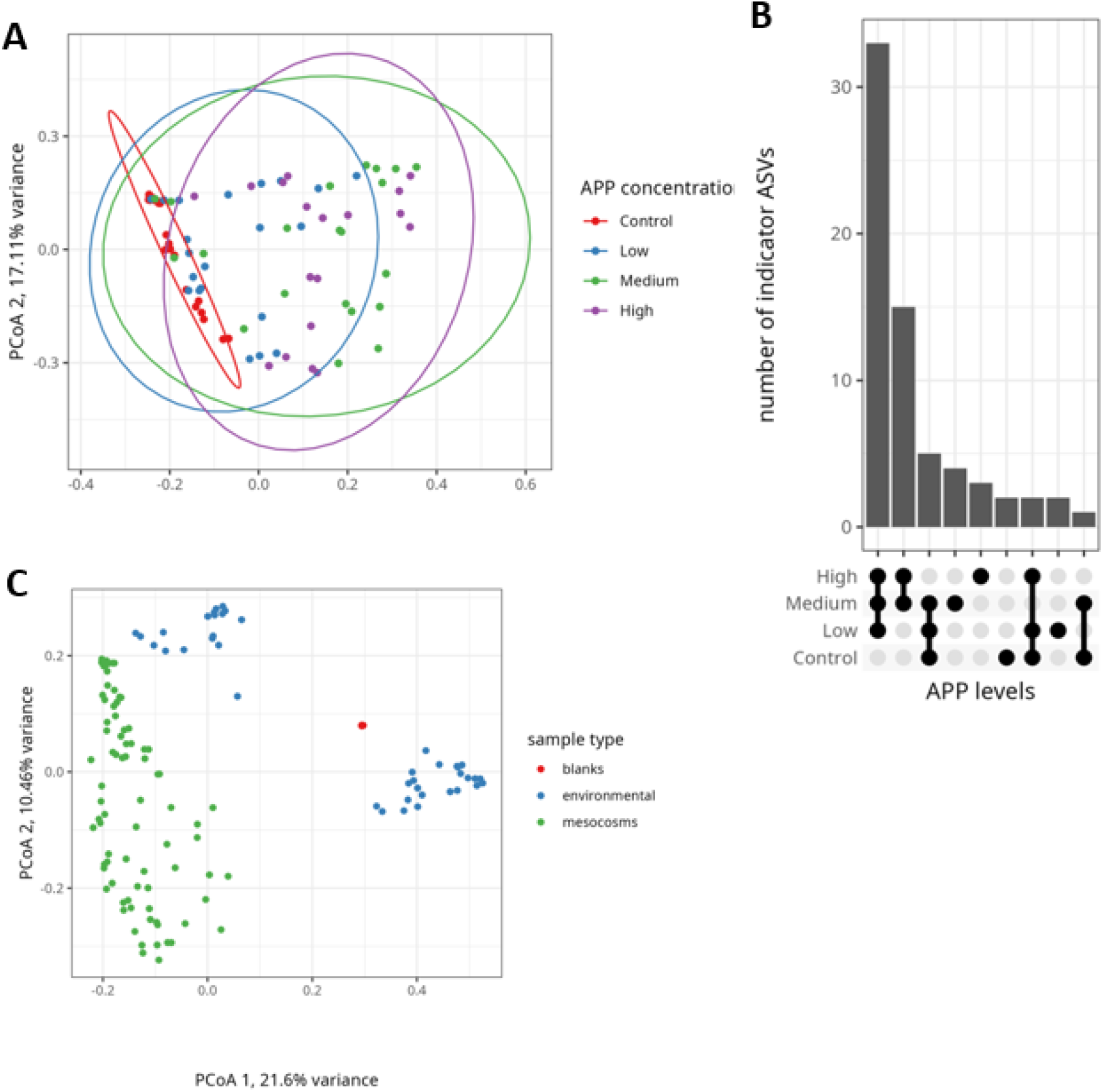
A: Principle coordinate analyses of 16S data based on Bray-Curtis dissimilarity matrix. Mesocosm samples distinguished by APP concentration. Control samples (no added APPs) denoted by red; low, medium and high APP concentrations denoted by blue, green and purple respectively. B: Results of indicator analysis detailing total indicators identified for each combination of sample groups. Contaminated (high, medium and low combined) vs non-contaminated (control) is marked by having by far the most indicators distinguishing the two groups. C: Principle coordinate analyses of 16S data based on Bray-Curtis dissimilarity matrix. Plot displays both mesocosm (green) and environmental samples (blue).

This finding is consistent with the results of an indicator analysis targeted at identifying indicators for the APP contamination levels, which results in the highest number of indicators distinguishing the APP-contaminated samples (considered collectively) from the control group (Figure 2B). Both the indicator analysis (full results in SI; Table S1) as well as a simple taxonomic analysis (see Figure 3) reveals ASVs belonging to the genus *Lutibacter* as relevant for the distinction between these two groups.

**Fig. 3.**
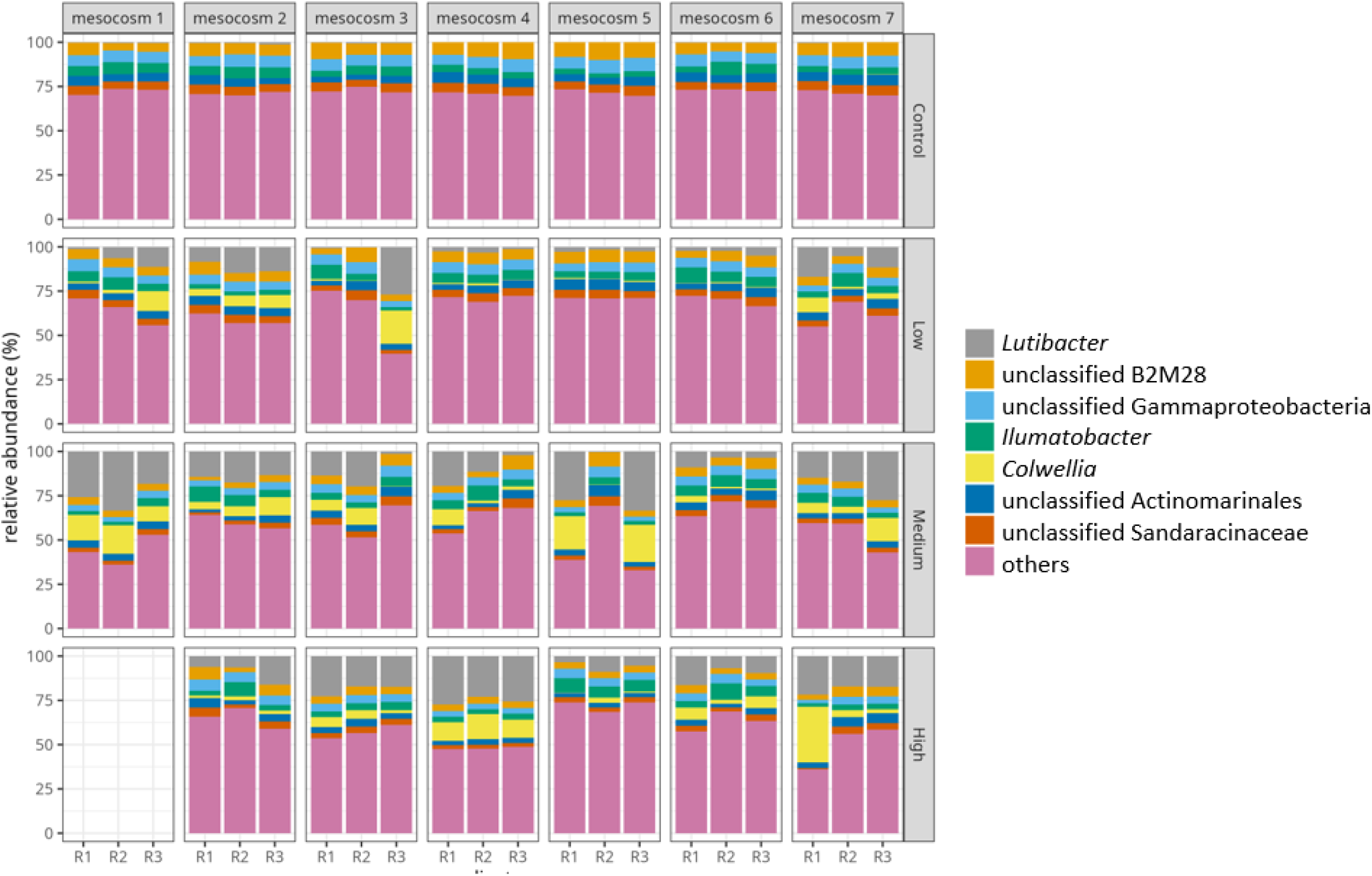
Barcharts visualising the top 7 genera by relative abundance across the mesocosm dataset. All other genera are combined in the “other” category. *Lutibacter*, the most prominent indicator genus for APP presence (as determined by indicator analysis) is shown in grey.

To determine if it would be possible to predict APP contamination levels from the 16S data, we trained and tested a Random Forest on the mesocosm data. While the model fails to reliably distinguish between APP contamination levels in testset samples with APP contamination (accuracy of 0.5; see Fig 4A), it almost perfectly distinguishes between samples with and without APP contamination (one false negative prediction, accuracy of ∼0.96; see Figure 4A)

**Fig. 4.**
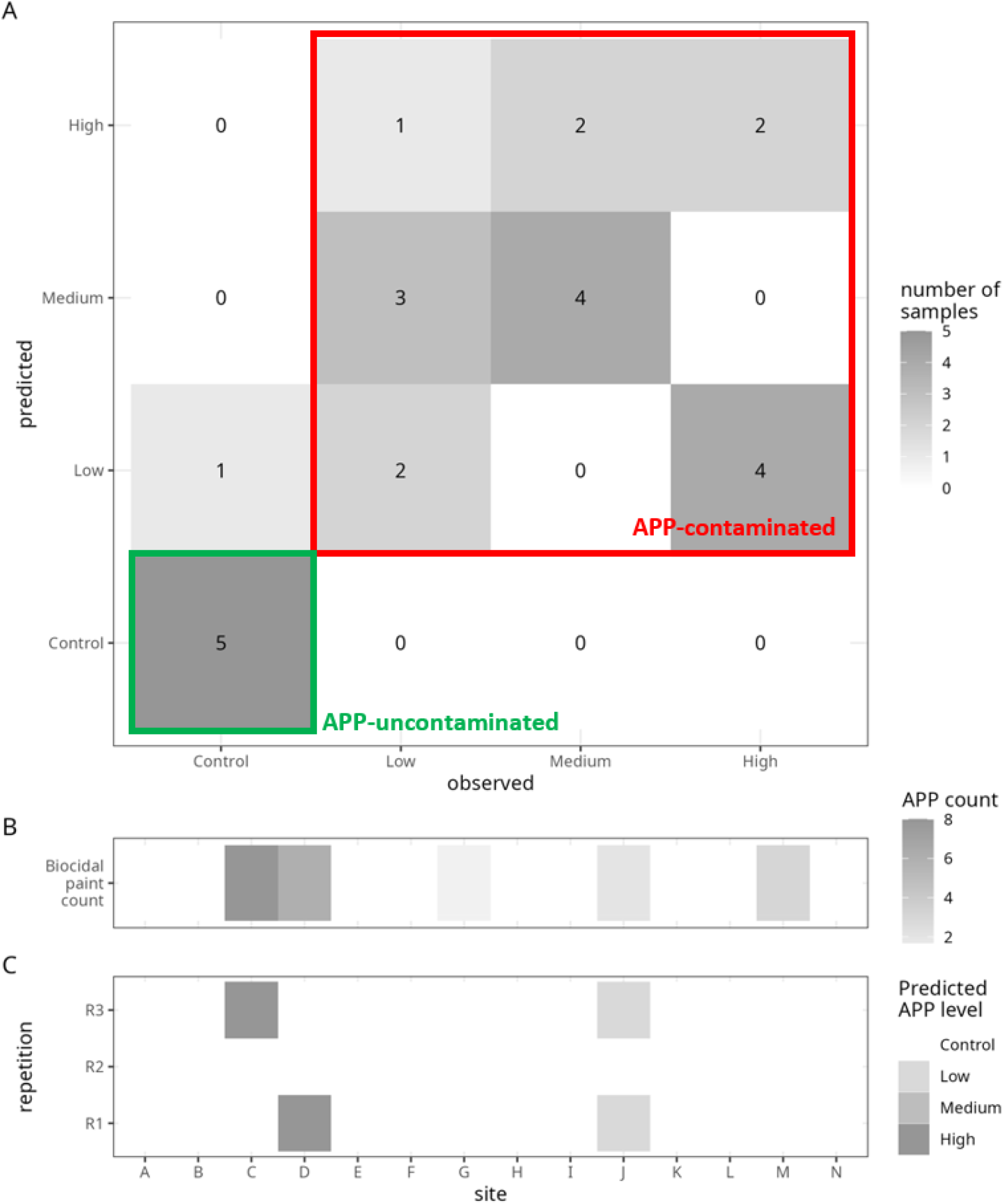
A: Confusion matrix showing model performance at predicting the testset data from the mesocosm experiment. While APP *concentration* is not well predicted, when low, medium and high are considered together as an overarching APP-contaminated group, prediction is highly successful with 83.35% correctly assigned APP-absence samples (5/6; green box) and 100% correctly assigned APP-presence samples (18/18; red box). B: APP counts determined in sediment from sites A - N using light microscopy-assisted visual selection with a Bogrov chamber combined with SEM-EDX spectroscopy. C: Model predictions for each environmental site based on the 16S rRNA sequencing data. Model correctly flags 3 of 5 contaminated sites, and correctly predicts all other sites as uncontaminated, giving an overall success rate of 85.7% (12/14).

### Contamination model extrapolates to environmental survey data – model validation

To validate the mesocosm experiment results with environmental samples, sediment samples were collected from 14 locations (Fig. 1B) and 85 particulates were visually identified as being possible paint particles and isolated for analytical chemistry characterisation (SEM-EDX). Of those 85 particulates, 20 were determined as APPs, across 5 different sites (see Figure 4B). Despite a large difference in microbial community composition between the samples from the mesocosm experiment and the environmental survey (Fig 2C) and differences in sediment type (Methods), the Random Forest model identified 3 sites as being APP-contaminated out of a total of 14 sites (see Figure 4C). All 3 of these sites were from verified APP-contaminated locations (see Figure 4B), thus 3 of the 5 contaminated sites were correctly flagged as contaminated by the model with no false-positive predictions. While the model did not identify all contaminated replicates within verified contaminated sites, this result is consistent with the expected spatial heterogeneity of APP distribution in natural sediments.

## Discussion

Antifouling paints are used as antibiotic agents on ship hulls; it is therefore to be somewhat expected that their presence in sediment has an effect on the composition of the microbial community. Here, we show that while this effect appears not to be markedly dose-dependent, at least within the range examined in this study, the presence of APPs induces a fundamental shift in microbial community composition. When viewed through this binary presence/absence lens, the random forest-based machine learning model trained and tested on the mesocosm experimental data and validated on the environmental sampling campaign data demonstrates clear viability. Remarkably, this model, trained on just 57 mesocosm samples from a single coastal site, correctly identified 3 contaminated sites from a spatial survey conducted across 14 different coastal and estuarine locations, without generating any false positives among the entire environmental dataset (Figure 4B-C). Machine learning approaches are known typically to rely on large datasets. Despite being trained on a relatively small dataset, the Random Forest model used here shows a strong predictive performance. This likely stems from the targeted nature of the experimental setup as opposed to training on real environmental samples which would be expected to have considerable variability simply based upon differing environmental factors at different locations. It is postulated that the mesocosm experiment minimized environmental variability unrelated to APP contamination while retaining real environmental conditions which can be lost in a laboratory-based microcosm approach, thereby allowing the model to focus more effectively on APP-specific microbial signals while retaining environmental relevance. Thus for the development of similar machine learning-based applications, especially when budgetary or other constraints restrict the ability to have many hundreds or thousands of samples sequenced, a targeted environment-based experimental setup could be the best approach to train and test a new model, especially at the exploratory or proof-of-concept phase.^22–24^

The predictive success of the model can be considered especially notable as the mesocosm experiment occurred between January - March, while sampling of the other sites did not begin until mid-April of the same year. As has been ubiquitously previously demonstrated, geographical location and time of year have a very strong influence on the microbial community composition.^25–28^ The substantial differences in community composition induced by spatial and temporal separation between the mesocosm and environmental samples are illustrated in Figure 2C, in which the mesocosm samples form a distinct cluster from the environmental samples, although the biggest distinction remains between estuarine and coastal sampling sites. Despite these differences, the Random Forest model was still able to identify a microbial “fingerprint” of APP contamination, suggesting that the structuring effect of APPs on the microbial community composition is very robust.

Furthermore, the APPs in the mesocosm experiment were freshly generated while the APPs in the field samples originate from degraded coatings. The ability of the Random Forest model to detect the latter while being trained on the former demonstrates that environmentally aged APPs still exert biologically relevant effects on microbial communities in their environment. That is to say, despite having aged to the point where particles have begun to be physically fragment from their original coating, these environmental APPs appear to still be leaching biocidal compounds at levels sufficient to alter microbial community composition. This finding supports earlier hypotheses made by the authors^7^ that the layered structure of antifouling coatings facilitate prolonged biocide leaching since as the film degrades a new coat of antifouling paint is continually exposed. However, these are the first results to demonstrate that environmental APPs, expected to be from aged and sufficiently degraded antifouling maritime coatings, exhibit comparable biological effects on microbial communities as newly manufactured particles.

Yet, while the model correctly identifies all 11 uncontaminated sites, and identifies 3 contaminated sites correctly, the remaining two contaminated sites were wrongly classified as uncontaminated. Furthermore, for all contaminated sites not all site replicates are flagged as contaminated. Indeed, for the most strongly contaminated sites (sites C and D; see figures 4B and 4C), only 1 of the 3 site replicates are correctly identified, and for the third remaining correctly-identified contaminated site (site J; see figures 4B and 4C), 2 of the 3 site replicates are identified as contaminated. These results reinforce the importance of adequate (minimum triplicate) replication in sediment microbial ecology studies. There are two plausible explanations for the inconsistent site replicate classification observed.

Firstly, the innate variability of microbial communities might obfuscate the contamination signal, particularly when APP concentration – or their biological effects – are low. This would explain the misclassification of site G, which exhibited the lowest APP abundance of all the contaminated sites. Secondly, the physical heterogeneity in the distribution of APP particles across the sediment at a given sampling site. At sites G and M, only one and two APPs, respectively, were identified across the three replicate sediment grabs. Thus, it should be expected that some replicates at low-level contamination sites would not contain APPs at all.

The indicator analysis allows for an insight into the microbial correlates of APP contamination; it determined a total of 31 ASVs as specifically APP-associated within the mesocosm experimental data (Fig 2C). Within this list of ASVs (see SI, Table S1), the genus *Lutibacter* stands out as representing 38.7% of the ASVs identified to be associated with APP contamination. Visualization of the taxonomic composition of mesocosm samples further supports the importance of *Lutibacter*, which is functionally absent from all control samples but prominent in most APP-contaminated samples (see Figure 3). However, drawing biologically-relevant conclusions as to why certain *Lutibacter* ASVs appear to profit from APP-contaminated conditions is constrained by the limited taxonomic resolution provided by the ∼290 bp long V4 section of 16S rRNA gene used for amplicon sequencing in this study.^29^ Further analyses using BLAST searches of the ASV sequences revealed no meaningful differences in the taxonomic make-up of the *Lutibacter* ASVs that have been identified as APP-contamination indicators (12 ASVs) and those that have not (10 ASVs; see SI Table S2); both groups largely matched to the type species *L. litoralis*, with both groups also having representation from *L. crassostreae, L. maritimus and L. ocean*i. However, it should be noted that such species-level assignments based on the short ∼290bp fragments are simply estimates which may well not be representative. So, while there appears to be no clear taxonomic distinctions between APP-indicative and non-APP-indicative *Lutibacter* ASVs based on these estimates, this may not be the case, and a more detailed look into this dynamic would be needed with longer read sequencing to fully determine this. And indeed, such an investigation could be interesting, since there are metabolic differences between different *Lutibacter* species. For example, based on type strain data^30^ starch can be degraded by *L. agarilyticus, L. litoralis and L. profundi* but not by *L. crassostreae, L. flavus, L. holmesii, L. maritimus, L. oricola* or *L. aestuarii* and casein can be degraded by *L. profundi and L. maritimus*, but cannot be degraded by any of the other aforementioned type strains. As such, further information on which specific species of *Lutibacter* show most preference for APP-contaminated conditions would be highly interesting in determining what metabolic implications APP pollution might instigate. One interesting aspect about some *Lutibacter* species (i.e. *L. aestuarii* and *L*. maritimus)^31,32^ is that they can be facultatively anaerobic, meaning that while they typically respire aerobically, they can also switch to fermentation under anoxic conditions. This could also be an interesting aspect to explain the better performance of certain *Lutibacter* ASVs under APP-affected conditions, since if APP-associated biocides cause changes to microbial communities which causes a shift to favour anaerobic/fermentation pathways, fast responding facultatively anaerobic taxa may benefit the most from such changing conditions. This study serves as a proof of concept that APP presence/absence can be detected and theoretically mapped using short-read (V4) 16S rRNA sequencing of microbial communities combined with Random Forest machine learning models. However, the development of a robust ecotoxicology monitoring tool applicable globally will require further refinement. For application in other coastal regions with distinct microbial communities and environmental conditions, additional model training and validation may be necessary, but pre-testing for model accuracy in other coastal regions, especially outside of the Baltic Sea, is required. Furthermore, the inclusion of more data with a more comparable contamination signal such as possibly provided by chemical analysis of sediment porewater for leached APP-associated factors, or at least copper content,^2^ might be a good strategy for future model expansion. However, to greatly increase the usefulness of such a model, expansion beyond APPs to include a larger range of anthropogenic stressors is encouraged. This study demonstrates the blueprint for how such a mesocosm experimental design can be used to train and test a predictive model for a key environmental pollutant, however predictive models for other pollutants including pharmaceuticals, especially antibiotics, heavy metals and pesticides could be designed, tested and validated in much a similar way. Even more indirect stressors which might appear alongside pollutants like nitrification or deoxygenation could also be incorporated using an adapted experimental setup as demonstrated within this study. Perhaps most crucial aspect in such an endeavour will be collaboration, since such stressors likely have considerable overlap, and microbial communities may well respond in similar, or compounding, ways. That APPs might act as a proxy for general anthropogenic stress is not a new idea; in a previous study by the authors it was demonstrated that the strongest APP indicators were generally much more well-described (taxonomically-classified) than the APP-absence indicators, suggesting hardiness to endure and succeed in APP-affected conditions may well also be the same hardiness which allows for laboratory cultivation, a precursor for microbial classification. Thus, if the effect of APPs is to simply force the most stress-resistant taxa to dominate it may well be that other anthropogenic stressors may force a similar community shift, meaning that perhaps anthropogenic stress generally might be predictable, but predicting presence of specific stressors where multiple might be at play is likely to be much more challenging. Indicator analysis will continue to be a crucial tool in determining how different anthropogenic stressors affect sediment microbial communities in ways both similar and different to APPs, especially to assist in the taxonomic explanation accompanied with the abstracted pattern recognition of machine learning tools.

In summary, using a specifically-designed mesocosm experimental setup, a predicative model was trained and tested on a relatively small number of samples (57 and 24 respectively) and demonstrated that APP-contaminated samples could be distinguished from uncontaminated samples based on 16S V4 rRNA sequencing data of the microbial community. When these results were validated using an environmental survey of 14 coastal and estuarine sites, the model correctly predicted 4 samples as APP-contaminated, across 3 sites. In total, APPs were found at 5 sites. While the model did not correctly predict every site replicate as contaminated where APPs were found, it is certain that some replicates at contaminated sites came from uncontaminated sediment grabs, so some replicates from an overall APP-positive site would be expected to be APP-negative. The model correctly predicted all uncontaminated samples, with zero false-positive predictions. Thus, even if the model still needs greater accuracy at predicting APP-contamination, its efficiency at determining low risk of APP-contamination is clear. This study stands as a proof-of-concept and a blueprint for a successful, small-scale experimental deign for the construction of machine learning models for anthropogenic stressor monitoring. More testing and data are needed, especially from sites beyond the analysed geographical area, for such a model to be scientifically viable as a fully-fledged ecotoxicology monitoring tool. Nevertheless, that the model works well for APP-contamination detection at a variety of sites, with very different environmental factors, sampled at a different time of year with distinctly different microbial community compositions, is arguably quite remarkable, making the future development of not only a complete APP monitoring tool, but a general anthropogenic stressor predictive tool very encouraging.

## Methods

### Mesocosm experiment

Chambers for the mesocosm experiment were constructed from polycarbonate housing. A stainless steel 304 mesh (ø = 120 µm) was used to encase the chambers to allow for water and microbial movement between the chamber and the surrounding sediment. A 1000 mm long stainless steel 304 rod was attached to each chamber to act as a sediment anchor and to enable the divers to find and retrieve the chambers easily. The cylindrical chambers had an internal volume of 274 cm^3^ (height = 80 mm, radius = 33 mm). APPs were produced by painting 2 layers of primer and 3 of antifouling top-coat on sheets of silicone rubber, before peeling partially cured paint away from the silicone rubber. The sheets of paint were then hung to fully cure before being milled and then sorted by sieving. Particles 500 – 1000 µm were used for chamber spiking. More information on the process of paint particle production can be found in Tagg^33^. 2 different types of APPs were used in a 50:50 ratio and manufacturer details of the primers and antifouling paints used can be found in the supplementary information (table S1). These were selected based on previous research by the authors investigating APP effects on sediment sin a microcosm, 1 APP type was selected as it had (marginally) the strongest effect on altering microbial communities compared to controls and the other was selected as it was the most analogous to all the other tested APPs (see Tagg et al.)^7^. 4 different concentrations were selected for testing: control (no paint), low (100 g m^3−1^), medium (250 g m^3−1^) and high (500 g m^3−1^). Based on the aforementioned volume of the chambers, this meant a mass of 0.27 g (low), 0.68 g (medium) and 1.37 g (high) were added to each appropriate chamber. For each concentration tested, 7 replicate chambers were deployed. APPs were sterilised by UV light exposure within a laminar flow cabinet for 15 minutes each side and then soaked in sterile (0.2 µm filtered) seawater for 14 days. Dried APPs were measured into appropriate masses and stored in air-tight sterile glass tubes until needed.

Chambers were deployed into sediment in January 2022 and collected in March 2022 after 60 days of exposure. Chambers were deployed at Heiligendamm in the Baltic sea, northern Germany, ∼ 100 m off the end of the Heiligendamm pier at a water depth of ∼10 m (54° 14’75.00” N 11° 84’25.59” E). Sediment was retrieved at the site by Van-Veen grab, the chambers were directly filled with sediment, APPs added, stirred to evenly distribute APPs through the sediment chamber and then placed into the sediment by divers, submerged into sediment so the lid of the chamber was at the sediment surface, thus the chambers were analogous to a sediment depth of 0 - 80 mm. Following exposure, the chambers were again retrieved by divers. 3 × 1g of sediment was immediately (on boat) extracted from each chamber and placed into 2 ml sterile centrifuge tubes (VWR, Germany). All tubes were flash frozen in liquid nitrogen and stored at −80°C awaiting DNA extraction. 1 chamber (high concentration) was lost due to the lid becoming removed during the 60 days of environmental exposure. Thus, a total of 27 chambers were retrieved, each which were sampled in triplicate giving a total of 81 samples from the mesocosm experiment.

### Validation survey sampling

Sediment samples were collected from both the Warnow estuary and local Baltic Sea coastline in NE Germany (see Figure 1 for sampling map). Samples were collected in April 2023. Sediment samples were collected using a Van-Veen sediment grab from either pier, outcrop or small sampling boat depending on the accessibility of the site. 3 grabs were taken at each site. For DNA extraction, just as for the aforementioned chamber study, 3 × 1g of sediment was collected and placed into 2 ml sterile centrifuge tubes (VWR, Germany). All tubes were stored on ice, until being flash frozen in liquid nitrogen and stored at −80°C. Sediment from all 3 grabs was homogenised and collected into 1 × 2L glass jars for paint particle presence (PP) analysis for each site.

### Validation survey paint particle analysis

Sediment from all 14 sites was analysed for potential PPs. It was determined, both as to be comparable to the manufactured and spiked APPs added to chambers, and for feasibility reasons (as already outlined in the introduction) to analyse for PPs >500 µm. Each 2L sediment sample was analysed identically. Sediment was firstly flushed through a 1 mm mesh-size sieve to remove any larger sediments particulates and then the effluent was then again passed through a 500 µm mesh-size sieve. All material collected on the sieve was then stored in covered sterile glass beakers. Each sample was then fluidised with ultra-high quality water and visually analysed, 5 ml at a time using a Bogorov chamber (Hydro-Bios Apparatebau GmbH, Germany) and a stereo microscope at 20 - 80 × magnification. Every particulate was visually assessed for PP potentiality according to standard protocols for visual assessment of MP.^34^ Every particle assessed to have any likelihood as being a PP was extracted from the Bogorov chamber with stainless steel tweezers and placed in a clean glass petri dish. Each particle was then analysed using SEM-EDX spectroscopy for antifouling-associated elemental profiles. SEM-EDX analyses were performed with a Zeiss Merlin Compact SEM (Zeiss, Germany) and Oxford Instruments EDX with Inca Feature 5:04 analysis software (Oxford Instruments, UK). Previous research by the author determined that of the wide array of commercially available biocidal antifouling paints available, all had discernable presence of copper (Cu) and/or zinc (Zn).^7^ Thus all fragments were categorised as either APP or non-APP based on if a given particle had a minimum of 1% Cu or Zn (weight %) in the surface chemistry. Other elemental indicators of now-banned biocidal chemistry such as tin (Sn) or arsenic (As) were also considered although no particles were detected to have such chemistry in any sample.

### DNA Extraction, Sequencing and Processing

DNA extraction was performed using Qiagen PowerSoil pro DNA extraction kits (Qiagen, USA) following manufacturer guidelines. Both extraction and PCR blanks were included. The V4 region of the 16S rRNA gene was selected for amplification using primers 515F (5′-GTGCCAGCMGCCGCGGTAA-3′) and 806R (5′ - GGACTACHVGGGTWTCTAAT - 3′).^35^ PCRs, library preparation and sequencing were performed according to the standard protocols for 16S metagenomic sequencing provided by Illumina.^36^

Primers were trimmed from demultiplexed sequencing reads using cutadapt version 4.2.^37^ Paired-end reads were then processed with dada2 version 1.26.0.^38^ to generate amplicon sequence variants (ASVs). Forward reads were trimmed to 150 bp, reverse to 140 bp, and filtered to a maximum expected error rate of 2. Performance of error learning was conducted with default settings. All samples were then pooled for denoising. Forward and reverse reads were merged with a minimum overlap of 10 bp. Chimera detection and removal was conducted using default settings. The *assignTaxonomy* function within dada2 using the SILVA ribosomal reference database version 138.1^39^ was used to taxonomically classify ASVs. The minimum bootstrap for taxon assignments of 70. Sequences affiliated with chloroplast and mitochondria were removed, along with any sequence which only appeared once in the dataset. Thus, only ASVs classified as bacteria or archaea with a sequence length between 251 bp and 257 bp were retained for the further analysis. All sequence processing steps were executed using snakemake workflow manager version 6.5.1.^40^

### Data Analysis

For all data analyses, OTU counts were turned into relative counts (via prop.table in R)^41^ for each sample and a rarity filter was applied. To this end, all OTUs were removed from the final 16S and 18S ASV tables that were not present in at least 1% of samples with 0.1% of relative abundance.Principal coordinate analysis (PCoA) ordinations were performed using the betadisper function on Bray-Curtis distance matrices calculated using the vegdist function (both from the package vegan, v2.6-6).^42^ Indicator species for the contaminations of APP levels were identified using the multipatt function from the indicspecies package (v 1.7.14)^43^ with 999 permutations.

Random forest models were trained to predict levels of APP contamination from community composition using the caret package (v 6.0-94, method = “rf”)^44^ on the OTU data from the mesocosm experiment using all replicates from two chambers of each level of APP contamination (C1, C7, C8, C14, C15, C21, C22, C27) as test dataset and the data from the others as training.

Visualizations were generated using ggplot2 (v 3.5.1),^45^ ggupset (v 0.4.0),^46^ patchwork (v 1.2.0)^47^ and the melt function from the reshape2 package (v 1.4.4).^48^ All code used in this study is available at https://git.io-warnemuende.de/bio_inf/IOWseq000057_PaintSed2. Sequencing data has been uploaded to NCBI and is under the following accession: PRJNA1304714.

## Acknowledgements

We would like to thank Janet Pissula for assistance with DNA extractions, Jana Bull for support with PCRs, library preparation and sequencing, and Daniel Herlemann and Christiane Hassenrück for their support with bioinformatics data processing and data curation. We would also like to thank Sascha Plewe for assistance with SEM-EDX analyses. This work was funded by the Deutsche Forschungsgemeinschaft (DFG) under grant number 452841212.

## Author Contributions

AT and ML conceived the original experimental ideas supported by TS; AT and AS undertook fieldwork; AT supervised & coordinated laboratory work; AS and AT performed laboratory work; TS undertook all data analysis and figure generation; AT wrote the first draft supported by TS; ML, AS and BK also contributed to the writing; AT secured funding for the research.

## References

1. Turner, A. Paint particles in the marine environment: An overlooked component of microplastics. Water Res. X 12, 100110 (2021).

2. Soroldoni, S. et al. Antifouling paint particles: Sources, occurrence, composition and dynamics. (2018) doi:10.1016/j.watres.2018.02.064.

3. Diana, Z. T., Chen, Y. & Rochman, C. M. Paint: a ubiquitous yet disregarded piece of the microplastics puzzle. Environ. Toxicol. Chem. 44, 26 (2025).

4. Which type of antifouling should I use on my boat? | International. https://www.international-yachtpaint.com/za/en/boat-paint-help/expert-advice/types-of-antifouling.

5. Kim, T., Eo, S., Shim, W. J. & Kim, M. Qualitative and quantitative assessment of microplastics derived from antifouling paint in effluent from ship hull hydroblasting and their emission into the marine environment. J. Hazard. Mater. 477, 135258 (2024).

6. Thomas, K. V., McHugh, M., Hilton, M. & Waldock, M. Increased persistence of antifouling paint biocides when associated with paint particles. Environ. Pollut. 123, 153–161 (2003).

7. Tagg, A. S. et al. Microplastic-antifouling paint particle contamination alters microbial communities in surrounding marine sediment. Sci. Total Environ. 926, 171863 (2024).

8. Turner, A., Ostle, C. & Wootton, M. Occurrence and chemical characteristics of microplastic paint flakes in the North Atlantic Ocean. Sci. Total Environ. 806, 150375 (2022).

9. Sparks, C. & Awe, A. Concentrations and risk assessment of metals and microplastics from antifouling paint particles in the coastal sediment of a marina in Simon’s Town, South Africa. Environ. Sci. Pollut. Res. 29, 59996–60011 (2022).

10. Kim, S. W., Song, W. Y., Waldman, W. R., Rillig, M. C. & Kim, T. Y. Toxicity of Aged Paint Particles to Soil Ecosystems: Insights from Caenorhabditis elegans. Environ. Sci. Technol. 58, 231–241 (2024).

11. Enders, K., Lenz, R., Ivar do Sul, J. A., Tagg, A. S. & Labrenz, M. When every particle matters: a QuEChERS approach to extract microplastics from environmental samples. MethodsX (2020) doi:10.1016/J.MEX.2020.100784.

12. Hidalgo-Ruz, V., Gutow, L., Thompson, R. C. & Thiel, M. Microplastics in the marine environment: A review of the methods used for identification and quantification. Environ. Sci. Technol. 46, 3060–75 (2012).

13. Sperlea, T., Heider, D. & Hattab, G. A theoretical basis for bioindication in complex ecosystems. Ecol. Indic. 140, (2022).

14. Zengel, S. et al. Impacts of the Deepwater Horizon Oil Spill on Salt Marsh Periwinkles (Littoraria irrorata). Environ. Sci. Technol. 50, 643–652 (2016).

15. Cordier, T., Lanzén, A., Apothéloz-Perret-Gentil, L., Stoeck, T. & Pawlowski, J. Embracing Environmental Genomics and Machine Learning for Routine Biomonitoring. Trends Microbiol. 27, 387–397 (2019).

16. Sperlea, T. et al. Quantification of the covariation of lake microbiomes and environmental variables using a machine learning-based framework. Mol. Ecol. 30, 2131–2144 (2021).

17. Hoshino, T. et al. Global diversity of microbial communities in marine sediment. Proc. Natl. Acad. Sci. U. S. A. 117, 27587–27597 (2020).

18. Ghannam, R. B. & Techtmann, S. M. Machine learning applications in microbial ecology, human microbiome studies, and environmental monitoring. Comput. Struct. Biotechnol. J. 19, 1092–1107 (2021).

19. Zschaubitz, E. et al. A benchmark analysis of feature selection and machine learning methods for environmental metabarcoding datasets. Comput. Struct. Biotechnol. J. 27, 1636–1647 (2025).

20. Janßen, R., Zabel, J., von Lukas, U. & Labrenz, M. An artificial neural network and Random Forest identify glyphosate-impacted brackish communities based on 16S rRNA amplicon MiSeq read counts. Mar. Pollut. Bull. 149, 110530 (2019).

21. Janßen, R. et al. A Glyphosate Pulse to Brackish Long-Term Microcosms Has a Greater Impact on the Microbial Diversity and Abundance of Planktonic Than of Biofilm Assemblages. Front. Mar. Sci. 6, 493759 (2019).

22. Janßen, R. et al. Machine Learning Predicts the Presence of 2,4,6-Trinitrotoluene in Sediments of a Baltic Sea Munitions Dumpsite Using Microbial Community Compositions. Front. Microbiol. 12, 626048 (2021).

23. Liu, B. et al. Machine learning-assisted identification of bioindicators predicts medium-chain carboxylate production performance of an anaerobic mixed culture. Microbiome 10, 1–21 (2022).

24. Loganathan, T. & Priya Doss C, G. The influence of machine learning technologies in gut microbiome research and cancer studies - A review. Life Sci. 311, 121118 (2022).

25. Jiang, P., Zhao, S., Zhu, L. & Li, D. Microplastic-associated bacterial assemblages in the intertidal zone of the Yangtze Estuary. Sci. Total Environ. 624, 48–54 (2018).

26. Wang, T. et al. Exploring changes in microplastic-associated bacterial communities with time, location, and polymer type in Liusha Bay, China. Mar. Environ. Res. 198, 106525 (2024).

27. Oberbeckmann, S., Kreikemeyer, B. & Labrenz, M. Environmental factors support the formation of specific bacterial assemblages on microplastics. Front. Microbiol. 8, 2709 (2018).

28. Oberbeckmann, S., Loeder, M. G. J., Gerdts, G. & Osborn, A. M. Spatial and seasonal variation in diversity and structure of microbial biofilms on marine plastics in Northern European waters. FEMS Microbiol. Ecol. 90, 478–492 (2014).

29. Buetas, E. et al. Full-length 16S rRNA gene sequencing by PacBio improves taxonomic resolution in human microbiome samples. BMC Genomics 25, 1–13 (2024).

30. Bauer, S. le M., Roalkvam, I., Steen, I. H. & Dahle, H. Lutibacter profundi sp. Nov., isolated from a deep-sea hydrothermal system on the arctic mid-ocean ridge and emended description of the genus lutibacter. Int. J. Syst. Evol. Microbiol. 66, 2671–2677 (2016).

31. Choi, D. H. & Cho, B. C. Lutibacter litoralis gen. nov., sp. nov., a marine bacterium of the family Flavobacteriaceae isolated from tidal flat sediment. Int. J. Syst. Evol. Microbiol. 56, 771–776 (2006).

32. Lee, S. Y., Lee, M. H., Oh, T. K. & Yoon, J. H. Lutibacter aestuarii sp. nov., isolated from a tidal flat sediment, and emended description of the genus Lutibacter Choi and Cho 2006. Int. J. Syst. Evol. Microbiol. 62, 420–424 (2012).

33. Tagg, A. S. Microplastic paint particle production for spiking experiments; silicone rubber as application material provide high yield with low effort. Microplastics Nanoplastics 2023 31 3, 1–5 (2023).

34. Gerdts, G. Standardised Protocol for Monitoring Microplastics in Sediments - Project BASEMANN Report. https://www.researchgate.net/publication/326552185 (2018) xdoi:10.13140/RG.2.2.36256.89601/1.

35. Caporaso, J. G. et al. Global patterns of 16S rRNA diversity at a depth of millions of sequences per sample. Proc. Natl. Acad. Sci. 108, 4516–4522 (2011).

36. Illumina. 16S metagenomic sequencing library preparation. Part # 15044223 Rev. B https://support.illumina.com/documents/documentation/chemistry_documentation/16s/16s-metagenomic-library-prep-guide-15044223-b.pdf.

37. Martin, M. Cutadapt removes adapter sequences from high-throughput sequencing reads. EMBnet.journal 17, 10–12 (2011).

38. Callahan, B. J. et al. DADA2: High-resolution sample inference from Illumina amplicon data. Nat. Methods 2016 137 13, 581–583 (2016).

39. Quast, C. et al. The SILVA ribosomal RNA gene database project: improved data processing and web-based tools. Nucleic Acids Res. 41, D590–D596 (2013).

40. Mölder, F. et al. Sustainable data analysis with Snakemake. F1000Research 2021 1033 10, 33 (2021).

41. R Core Team. R: A language and environment for statistical computing. at https://www.r-project.org (2022).

42. Oksanen, J. et al. Vegan: community ecology package. R package version 2.0-10. (2013).

43. De Cáceres, M. & Legendre, P. Associations between species and groups of sites: indices and statistical inference. Ecology 90, 3566–3574 (2009).

44. Kuhn, M. Building Predictive Models in R Using the caret Package. J. Stat. Softw. 28, 1–26 (2008).

45. Wickham, H. Ggplot2: Elegant Graphics for Data Analysis. (Springer-Verlag, New York, US, 2016).

46. Ahlmann-Eltze, C. Combination Matrix Axis for ‘ggplot2’ to Create ‘UpSet’ Plots [R package ggupset version 0.3.0]. (2020).

47. Pedersen, T. L. The Composer of Plots [R package patchwork version 1.3.1]. CRAN Contrib. Packag. (2025) doi:10.32614/CRAN.PACKAGE.PATCHWORK.

48. reshape2: Flexibly Reshape Data: A Reboot of the Reshape Package. Hadley Wickham (2020) doi:10.32614/CRAN.PACKAGE.RESHAPE2.

