## Supplementary material for "Predicting Antifouling Paint Particle Contamination based on 16S rRNA Gene Sequencing Data using Random Forest-Based Machine Learning": SI

### Supplementary Information

**Table S1.** Indicator analysis results between combinations of sample types. Index refers to the numeric ID for different combinations of sample group comparisons. Index 14 refers to APP-presence indicators (i.e. low, medium and high vs control). Kingdom column removed since every taxonomy in the table belongs to Bacteria. Statistic refers to the association/fidelity statistic and p.value refers to the P value of the permutation test.

| ASV | Control | High | Low | Medium | Index | Statistic | p.value | phylum | class | order | family | genus |
|---|---|---|---|---|---|---|---|---|---|---|---|---|
| ASV 6 | 0 | 1 | 1 | 1 | 14 | 0.999 | 0.001 | Proteobacteria | Gammaproteobacteria | Enterobacterales | Colwelliaceae | *Colwellia* |
| ASV 7 | 0 | 1 | 1 | 1 | 14 | 0.999 | 0.001 | Bacteroidota | Bacteroidia | Flavobacteriales | Flavobacteriaceae | *Lutibacter* |
| ASV 164 | 0 | 1 | 1 | 1 | 14 | 0.997 | 0.001 | Bacteroidota | Bacteroidia | Bacteroidales | Marinifilaceae | *Marinifilum* |
| ASV 122 | 0 | 1 | 1 | 1 | 14 | 0.991 | 0.001 | Bacteroidota | Bacteroidia | Flavobacteriales | Flavobacteriaceae | *Lutibacter* |
| ASV 554 | 0 | 1 | 1 | 1 | 14 | 0.975 | 0.001 | Proteobacteria | Gammaproteobacteria | Enterobacterales | Colwelliaceae | *Colwellia* |
| ASV 563 | 0 | 1 | 1 | 1 | 14 | 0.974 | 0.001 | Bacteroidota | Bacteroidia | Flavobacteriales | Flavobacteriaceae | *Lutibacter* |
| ASV 518 | 0 | 1 | 1 | 1 | 14 | 0.972 | 0.001 | Bacteroidota | Bacteroidia | Flavobacteriales | Flavobacteriaceae | *Lutibacter* |
| ASV 220 | 0 | 1 | 1 | 1 | 14 | 0.965 | 0.001 | Firmicutes | Clostridia | Peptostreptococcales-Tissierellales | Fusibacteraceae | *Fusibacter* |
| ASV 124 | 0 | 1 | 1 | 1 | 14 | 0.962 | 0.001 | Proteobacteria | Gammaproteobacteria | Pseudomonadales | Nitrincolaceae | *Motiliproteus* |

Table S1 continued

| | | | | | | | | | | | | |
|---|---|---|---|---|---|---|---|---|---|---|---|---|
| ASV 1085 | 0 | 1 | 1 | 1 | 14 | 0.931 | 0.001 | Firmicutes | Clostridia | Lachnospirales | Lachnospiraceae | Lachnospiraceae unclassified |
| ASV 291 | 0 | 1 | 1 | 1 | 14 | 0.913 | 0.001 | Bacteroidota | Bacteroidia | Flavobacteriales | Flavobacteriaceae | *Lutibacter* |
| ASV 540 | 0 | 1 | 1 | 1 | 14 | 0.913 | 0.001 | Proteobacteria | Gammaproteobacteria | Enterobacterales | Colwelliaceae | *Colwellia* |
| ASV 1017 | 0 | 1 | 1 | 1 | 14 | 0.904 | 0.001 | Bacteroidota | Bacteroidia | Flavobacteriales | Flavobacteriaceae | *Lutibacter* |
| ASV 1544 | 0 | 1 | 1 | 1 | 14 | 0.893 | 0.001 | Bacteroidota | Bacteroidia | Flavobacteriales | Flavobacteriaceae | *Lutibacter* |
| ASV 830 | 0 | 1 | 1 | 1 | 14 | 0.876 | 0.047 | Proteobacteria | Gammaproteobacteria | Enterobacterales | Alteromonadaceae | *Paraglaciecola* |
| ASV 476 | 0 | 1 | 1 | 1 | 14 | 0.873 | 0.009 | Bacteroidota | Bacteroidia | Flavobacteriales | Flavobacteriaceae | *Lutibacter* |
| ASV 1437 | 0 | 1 | 1 | 1 | 14 | 0.865 | 0.002 | Desulfobacterota | Desulfuromonadia | Desulfuromonadales | [illegible]eae | *Desulfuromusa* |
| ASV 1625 | 0 | 1 | 1 | 1 | 14 | 0.865 | 0.001 | Firmicutes | Clostridia | Lachnospirales | Lachnospiralesunclassified | Lachnospirales unclassified |
| ASV 1483 | 0 | 1 | 1 | 1 | 14 | 0.862 | 0.018 | Proteobacteria | Gammaproteobacteria | Enterobacterales | Shewanellaceae | *Shewanella* |
| ASV 1779 | 0 | 1 | 1 | 1 | 14 | 0.856 | 0.001 | Bacteroidota | Bacteroidia | Flavobacteriales | Flavobacteriaceae | *Lutibacter* |
| ASV 562 | 0 | 1 | 1 | 1 | 14 | 0.837 | 0.001 | Proteobacteria | Gammaproteobacteria | Methylococcales | Cycloclasticaceae | *Cycloclasticus* |
| ASV 5907 | 0 | 1 | 1 | 1 | 14 | 0.827 | 0.001 | Proteobacteria | Gammaproteobacteria | Pseudomonadales | Pseudomonadalesunclassified | Pseudomonadales unclassified |
| ASV 2169 | 0 | 1 | 1 | 1 | 14 | 0.823 | 0.001 | Proteobacteria | Gammaproteobacteria | Enterobacterales | Kangiellaceae | *Aliikangiella* |
| ASV 912 | 0 | 1 | 1 | 1 | 14 | 0.816 | 0.001 | Bacteroidota | Bacteroidia | Flavobacteriales | Flavobacteriaceae | *Lutibacter* |
| ASV 944 | 0 | 1 | 1 | 1 | 14 | 0.816 | 0.001 | Bacteroidota | Bacteroidia | Flavobacteriales | Flavobacteriaceae | *Lutibacter* |
| ASV 1268 | 0 | 1 | 1 | 1 | 14 | 0.815 | 0.001 | Proteobacteria | Gammaproteobacteria | Thiotrichales | Thiotrichaceae | Thiotrichaceae unclassified |
| ASV 583 | 0 | 1 | 1 | 1 | 14 | 0.806 | 0.001 | Patescibacteria | Gracilibacteria | JGI0000069-P22 | JGI0000069-P22unclassified | JGI0000069-P22 unclassified |
| ASV 3243 | 0 | 1 | 1 | 1 | 14 | 0.777 | 0.001 | Bacteroidota | Bacteroidia | Flavobacteriales | Flavobacteriaceae | *Lutibacter* |
| ASV 1071 | 0 | 1 | 1 | 1 | 14 | 0.766 | 0.012 | Proteobacteria | Gammaproteobacteria | Enterobacterales | Psychromonadaceae | *Psychromonas* |
| ASV 2702 | 0 | 1 | 1 | 1 | 14 | 0.606 | 0.006 | Proteobacteria | Gammaproteobacteria | Methylococcales | Cycloclasticaceae | *Cycloclasticus* |
| ASV 2105 | 0 | 1 | 1 | 1 | 14 | 0.582 | 0.020 | Bacteroidota | Bacteroidia | Flavobacteriales | Flavobacteriaceae | *Flavobacterium* |

Table S1 continued

| | | | | | | | | | | | | |
|---|---|---|---|---|---|---|---|---|---|---|---|---|
| ASV 2095 | 1 | 0 | 1 | 1 | 13 | 0.795 | 0.015 | Proteobacteria | Gammaproteobacteria | Steroidobacterales | Woeseiaceae | *Woeseia* |
| ASV 1343 | 1 | 0 | 1 | 1 | 13 | 0.792 | 0.005 | Cyanobacteria | Cyanobacteriia | Leptolyngbyales | Leptolyngbyaceae | CalothrixKVSF5 |
| ASV 1145 | 1 | 0 | 1 | 1 | 13 | 0.749 | 0.003 | Actinobacteriota | Acidimicrobiia | Microtrichales | Microtrichaceae | Microtrichaceae unclassified |
| ASV 776 | 1 | 0 | 1 | 1 | 13 | 0.736 | 0.010 | Bacteroidota | Bacteroidia | Flavobacteriales | Flavobacteriaceae | Flavobacteriaceae unclassified |
| ASV 1029 | 1 | 0 | 1 | 1 | 13 | 0.724 | 0.013 | Sva0485 | Sva0485 unclassified | Sva0485 unclassified | unclassified | unclassified |
| ASV 1250 | 1 | 0 | 1 | 1 | 13 | 0.710 | 0.015 | Planctomycetota | Planctomycetes | Planctomycetales | Rubinisphaeraceae | Rubinisphaeracea e unclassified |
| ASV 1323 | 1 | 0 | 1 | 1 | 13 | 0.704 | 0.018 | Actinobacteriota | Acidimicrobiia | Microtrichales | Microtrichaceae | Microtrichaceae unclassified |
| ASV 1058 | 1 | 0 | 1 | 1 | 13 | 0.681 | 0.007 | Planctomycetota | Planctomycetes | Pirellulales | Pirellulaceae | *Rubripirellula* |
| ASV 1670 | 1 | 0 | 1 | 1 | 13 | 0.676 | 0.043 | Gemmatimonadota | BD2-11 terrestrial group | BD2-11 terrestrial group unclassified | 11terrestrialgroup unclassified | BD2-11 terrestrial group unclassified |
| ASV 1042 | 1 | 0 | 1 | 1 | 13 | 0.658 | 0.048 | Bacteroidota | Bacteroidia | Flavobacteriales | Flavobacteriaceae | Flavobacteriaceae unclassified |
| ASV 2257 | 1 | 0 | 1 | 1 | 13 | 0.653 | 0.028 | Proteobacteria | Gammaproteobacteria | BD7-8 | BD7-8 unclassified | BD7-8 unclassified |
| ASV 962 | 1 | 1 | 1 | 0 | 11 | 0.954 | 0.001 | LCP-89 | LCP-89 unclassified | LCP-89 unclassified | LCP-89 unclassified | unclassified |
| ASV 2694 | 1 | 1 | 1 | 0 | 11 | 0.761 | 0.007 | Desulfobacterota | Desulfuromonadia | Desulfuromonadales | ae | *Trichloromonas* |
| ASV 1055 | 1 | 1 | 1 | 0 | 11 | 0.748 | 0.008 | Desulfobacterota | Desulfobacteria | Desulfobacterales | Desulfobacteraceae | *Desulfobacula* |
| ASV 1826 | 1 | 1 | 1 | 0 | 11 | 0.610 | 0.042 | Bacteroidota | Bacteroidia | Bacteroidales | Marinilabiliaceae | *Carboxylicivirga* |
| ASV 1417 | 0 | 0 | 1 | 1 | 10 | 0.554 | 0.019 | Bacteroidota | Bacteroidia | BacteroidetesVC2.1Ba c22 | BacteroidetesVC2.1B ac22 unclassified | Bacteroidetes VC2.1Bac22 unclassified |
| ASV 677 | 0 | 1 | 0 | 1 | 9 | 0.960 | 0.001 | Proteobacteria | Gammaproteobacteria | Pseudomonadales | Spongiibacteraceae | *Dasania* |
| ASV 894 | 0 | 1 | 0 | 1 | 9 | 0.916 | 0.001 | Proteobacteria | Alphaproteobacteria | Rhodobacterales | Rhodobacteraceae | Rhodobacteraceae unclassified |
| ASV 1587 | 0 | 1 | 0 | 1 | 9 | 0.874 | 0.001 | Proteobacteria | Gammaproteobacteria | Enterobacterales | Colwelliaceae | *Colwellia* |
| ASV 605 | 0 | 1 | 0 | 1 | 9 | 0.856 | 0.001 | Proteobacteria | Gammaproteobacteria | Enterobacterales | Colwelliaceae | *Colwellia* |
| ASV 991 | 0 | 1 | 0 | 1 | 9 | 0.846 | 0.001 | Proteobacteria | Gammaproteobacteria | Pseudomonadales | Nitrincolaceae | *Motiliproteus* |

Table S1 continued

| | | | | | | | | | | | | |
|---|---|---|---|---|---|---|---|---|---|---|---|---|
| ASV 950 | 0 | 1 | 0 | 1 | 9 | 0.766 | 0.001 | Proteobacteria | Alphaproteobacteria | Rhodobacterales | Rhodobacteraceae | Rhodobacteraceae unclassified |
| ASV 1876 | 0 | 1 | 0 | 1 | 9 | 0.748 | 0.002 | Proteobacteria | Gammaproteobacteria | Enterobacterales | Psychromonadaceae | *Psychromonas* |
| ASV 3162 | 0 | 1 | 0 | 1 | 9 | 0.742 | 0.003 | Proteobacteria | Gammaproteobacteria | Pseudomonadales | Pseudomonadaceae | *Pseudomonas* |
| ASV 1567 | 0 | 1 | 0 | 1 | 9 | 0.716 | 0.003 | Campylobacterota | Campylobacteria | Campylobacterales | Arcobacteraceae | *Pseudarcobacter* |
| ASV 2182 | 0 | 1 | 0 | 1 | 9 | 0.712 | 0.033 | Bacteroidota | Bacteroidia | Flavobacteriales | Flavobacteriaceae | *Cellulophaga* |
| ASV 1576 | 0 | 1 | 0 | 1 | 9 | 0.619 | 0.002 | Proteobacteria | Gammaproteobacteria | Pseudomonadales | Pseudomonadaceae | *Pseudomonas* |
| ASV 4484 | 0 | 1 | 0 | 1 | 9 | 0.595 | 0.010 | Proteobacteria | Gammaproteobacteria | Pseudomonadales | Pseudomonadaceae | *Pseudomonas* |
| ASV 2418 | 0 | 1 | 0 | 1 | 9 | 0.584 | 0.003 | Proteobacteria | Gammaproteobacteria | Enterobacterales | Colwelliaceae | *Colwellia* |
| ASV 4139 | 1 | 0 | 0 | 1 | 7 | 0.510 | 0.034 | Bacteroidota | Bacteroidia | Bacteroidales | Bacteroidales unclassified | Bacteroidales unclassified |
| ASV 2272 | 1 | 0 | 0 | 1 | 7 | 0.443 | 0.030 | Acidobacteriota | Blastocatellia | Blastocatellales | Blastocatellaceae | *Blastocatella* |
| ASV 1221 | 1 | 0 | 1 | 0 | 6 | 0.763 | 0.001 | Planctomycetota | Planctomycetes | Pirellulales | Pirellulaceae | *Rhodopirellula* |
| ASV 726 | 1 | 0 | 1 | 0 | 6 | 0.753 | 0.001 | Bacteroidota | Bacteroidia | Bacteroidales | Prolixibacteraceae | *Draconibacterium* |
| ASV 1228 | 1 | 0 | 1 | 0 | 6 | 0.632 | 0.008 | Nanoarchaeota | Nanoarchaeia | Woesearchaeales | Woesearchaeales unclassified | Woesearchaeales unclassified |
| ASV 1467 | 1 | 0 | 1 | 0 | 6 | 0.550 | 0.004 | Proteobacteria | Gammaproteobacteria | Ectothiorhodospirales | eae | *Thiogranum* |
| ASV 619 | 1 | 0 | 1 | 0 | 6 | 0.547 | 0.006 | Cyanobacteria | Cyanobacteriia | Cyanobacteriales | Xenococcaceae | Pleurocapsa PCC-7319 |
| ASV 2434 | 0 | 0 | 0 | 1 | 4 | 0.803 | 0.001 | Proteobacteria | Gammaproteobacteria | Enterobacterales | Colwelliaceae | *Colwellia* |
| ASV 3154 | 0 | 0 | 0 | 1 | 4 | 0.484 | 0.005 | Bacteroidota | Bacteroidia | Flavobacteriales | Flavobacteriaceae | *Lutibacter* |
| ASV 3783 | 0 | 0 | 1 | 0 | 3 | 0.699 | 0.001 | Bacteroidota | Bacteroidia | Bacteroidales | Marinifilaceae | Marinifilaceae unclassified |
| ASV 187 | 0 | 0 | 1 | 0 | 3 | 0.553 | 0.001 | Bacteroidota | Bacteroidia | Bacteroidales | Bacteroidales unclassified | Bacteroidales unclassified |
| ASV 5083 | 0 | 1 | 0 | 0 | 2 | 0.705 | 0.001 | Firmicutes | Clostridia | Peptostreptococcales-Tissierellales | Fusibacteraceae | *Fusibacter* |
| ASV 939 | 0 | 1 | 0 | 0 | 2 | 0.408 | 0.005 | Campylobacterota | Campylobacteria | Campylobacterales | Sulfurovaceae | *Sulfurovum* |
| ASV 1628 | 1 | 0 | 0 | 0 | 1 | 0.559 | 0.020 | Desulfobacterota | Desulfobulbia | Desulfobulbales | Desulfocapsaceae | Desulfocapsaceae unclassified |
| ASV 1176 | 1 | 0 | 0 | 0 | 1 | 0.415 | 0.024 | Actinobacteriota | Acidimicrobiia | Microtrichales | Microtrichaceae | Microtrichaceae unclassified |

**Table S2.** NCBI BLAST best hits for all *Lutibacter* ASVs, organised by APP indicator status.

| Indicator ASVs | *NCBI* accession | Taxonomy | Percent Identity (%) | Query Coverage |
|---|---|---|---|---|
| ASV 7 | gi\|1477912639\|gb\|MH929593.1\| | *Lutibacter litoralis* | 99.209 | 100 |
| ASV 122 | gi\|1756308240\|gb\|MN535761.1\| | *Lutibacter agarilyticus* | 99.605 | 100 |
| ASV 563 | gi\|1477912639\|gb\|MH929593.1\| | *Lutibacter litoralis* | 98.814 | 100 |
| ASV 518 | gi\|1477912639\|gb\|MH929593.1\| | *Lutibacter litoralis* | 99.605 | 100 |
| ASV 291 | gi\|1477912639\|gb\|MH929593.1\| | *Lutibacter litoralis* | 98.814 | 100 |
| ASV 1017 | gi\|1024974821\|ref\|NR_136818.1\| | *Lutibacter crassostreae* | 99.209 | 100 |
| ASV 1544 | gi\|1779258078\|gb\|MN746248.1\| | *Lutibacter maritimus* | 99.605 | 100 |
| ASV 476 | gi\|1477912639\|gb\|MH929593.1\| | *Lutibacter litoralis* | 99.209 | 100 |
| ASV 1779 | gi\|2155011107\|gb\|OL629130.1\| | *Lutibacter maritimus* | 98.024 | 100 |
| ASV 912 | gi\|1477912639\|gb\|MH929593.1\| | *Lutibacter litoralis* | 98.814 | 100 |
| ASV 944 | gi\|1477912639\|gb\|MH929593.1\| | *Lutibacter litoralis* | 99.605 | 100 |
| ASV 3243 | gi\|1182956133\|ref\|NR_146841.1\| | *Lutibacter oceani* | 98.419 | 100 |
| **Non-indicative Lutibacter ASVs** | | | | |
| ASV 5 | gi\|1477912639\|gb\|MH929593.1\| | *Lutibacter litoralis* | 100 | 100 |
| ASV 217 | gi\|1477912639\|gb\|MH929593.1\| | *Lutibacter litoralis* | 99.209 | 100 |
| ASV 279 | gi\|1024974821\|ref\|NR_136818.1\| | *Lutibacter crassostreae* | 98.814 | 100 |
| ASV 408 | gi\|1182956133\|ref\|NR_146841.1\| | *Lutibacter oceani* | 98.814 | 100 |
| ASV 692 | gi\|1779258078\|gb\|MN746248.1\| | *Lutibacter maritimus* | 98.024 | 100 |
| ASV 713 | gi\|2155011107\|gb\|OL629130.1\| | *Lutibacter maritimus* | 99.605 | 100 |
| ASV 1110 | gi\|1477912639\|gb\|MH929593.1\| | *Lutibacter litoralis* | 98.814 | 100 |
| ASV 2082 | gi\|1024974821\|ref\|NR_136818.1\| | *Lutibacter crassostreae* | 99.605 | 100 |
| ASV 2332 | gi\|374431280\|gb\|JQ241143.1\| | *Lutibacter holmesii* | 98.814 | 100 |
| ASV 3154 | gi\|1477912639\|gb\|MH929593.1\| | *Lutibacter litoralis* | 99.605 | 100 |
